# Draft genome of the Reindeer (*Rangifer tarandus*)

**DOI:** 10.1101/154724

**Authors:** Zhipeng Li, Zeshan Lin, Lei Chen, Hengxing Ba, Yongzhi Yang, Kun Wang, Wen Wang, Qiu Qiang, Guangyu Li

## Abstract

**Background:** Reindeer (*Rangifer tarandus*) is the only fully domesticated species in the Cervidae family, and is the only cervid with a circumpolar distribution. Unlike all other cervids, female reindeer regularly grow cranial appendages (antlers, the defining characteristics of cervids), as well as males. Moreover, reindeer milk contains more protein and less lactose than bovids’ milk. A high quality reference genome of this specie will assist efforts to elucidate these and other important features in the reindeer.

**Findings:** We obtained 723.2 Gb (Gigabase) of raw reads by an Illumina Hiseq 4000 platform, and a 2.64 Gb final assembly, representing 95.7% of the estimated genome (2.76 Gb according to k-mer analysis), including 92.6% of expected genes according to BUSCO analysis. The contig N50 and scaffold N50 sizes were 89.7 kilo base (kb) and 0.94 mega base (Mb), respectively. We annotated 21,555 protein-coding genes and 1.07 Gb of repetitive sequences by *de novo* and homology-based prediction. Homology-based searches detected 159 rRNA, 547 miRNA, 1,339 snRNA and 863 tRNA sequences in the genome of *R. tarandus*. The divergence time between *R. tarandus*, and ancestors of *Bos taurus* and *Capra hircus*, is estimated to be 29.55 million years ago (Mya).

**Conclusions:** Our results provide the first high-quality reference genome for the reindeer, and a valuable resource for studying evolution, domestication and other unusual characteristics of the reindeer.

## Background information

The Cervidae is the second largest family in the suborder Ruminantia of the Artiodactyla, which are distributed across much of the globe in diverse habitats, from arctic tundra to tropical forests [1, 2]. Interestingly, reindeer (*Rangifer tarandus*) is the only species with a circumpolar distribution (present in boreal, tundra, subarctic, arctic and mountainous regions of northern Asia, North America and Europe). It is also the only cervid having been fully domesticated, although some other species, such as the sika deer (*Cervus nippon*), which has been semi-domesticated for more than 200 years and still has strong wild nature. Antlers, male secondary sexual appendage, are the defining characteristic of cervids, which shed and regrow each year throughout an animal’s life. However, reindeer do not follow this rule, with the exception in which females also bear shedding antlers. Moreover, reindeer milk contains greater amount of proteins, and lower amount of lactose compared to that of bovids [3]. Here, we report a high-quality reindeer reference genome using material from a Chinese individual, which will be useful in elucidating special characteristics of special cervid.

## Data description

### Animal and sample collecting

Fresh blood was collected from a two-year-old, female reindeer of a domesticated herd maintained by Ewenki hunter-herders in the Greater Khingan Mountains, Inner Mongolia Autonomous Region, China (50.77° N, 121.47° E). The sample was immediately placed in liquid nitrogen, and was then stored at -80°C for later analysis.

### Library construction, sequencing and filtering

Genomic DNA was extracted from the fresh blood. The isolated genomic DNA was then used to construct five short-insert libraries (200, 250, 350, 400 and 450 base pair, bp) and four long-insert libraries (3, 6.5, 11.5 and 16 kb) following standard protocols provided by Illumina. Then, 150 bp paired-end sequencing was performed to generate 723.2 Gb raw data, using a whole genome shotgun sequencing strategy on an Illumina Hiseq 4000 platform (**Table S1**). To improve the quality of reads, we trimmed low-quality bases from both sides of reads and removed reads with more than 5% of uncalled (“N”) bases. Then reads of all libraries were corrected by SOAPec (version 2.03) [4]. Finally, clean reads amounting to 615 Gb were obtained for genome assembly.

### Evaluation of genome size

The estimated genome size is 2.76 Gb according to k-mer analysis, based on the following formula: G = k-mer_number/k-mer_depth (**Figure S1**) [5]. All the clean reads provide approximately ~ 220-fold mean coverage.

### Genome assembly

We used SOAPdenovo (version 2.04) with optimized parameters (pregraph −K 79 −d 0; map -k 79; scaff -L 200) to construct contigs and original scaffolds [5]. All reads were aligned onto contigs for scaffold construction by utilizing the paired-end information. Gaps were filled using reads from three libraries (200, 250 and 350 bp) with GapCloser (version 1.12) [6]. The final reindeer genome assembly is 2.64 Gb long, including 95.7 Mb (3.6%) of unknown bases, smaller than that of the domestic goat (*Capra hircus*, 2.92 Gb) [7], and similar to that of sheep (*Ovis aries*, 2.61 Gb) [8]. The contig N50 (> 200 bp) and scaffold N50 (> 500 bp) sizes are 89.7 kb and 0.94 Mb, respectively (**Table 1**).

**Table 1.** Summary of the genome assembly of *Rangier tarandus*.

| Type | Scaffold (bp) | Contig (bp) |
| --- | --- | --- |
| Total number | 58,765 | 117,102 |
| Total length | 2,832,785,815 | 2,732,476,387 |
| N50 length | 986,392 | 91,805 |
| N90 length | 151,297 | 17,480 |
| Max length | 4,664,725 | 770,474 |
| GC content(%) | 41.24 | 40.98 |

### Quality assessments of the assembled genome

We used BUSCO (benchmarking universal single-copy orthologs, version 2.0) software to assess the genome completeness [9]. Our assembly covered 92.6% of the core genes, with 3,803 genes being complete (**Table S2**). Feature-response curve (FRC, version 1.3.1) method [10] was then used to evaluate the trade-off between the assembly’s contiguity and correctness. The results indicate that it has similar accumulated curve compare to published high quality assemblies for ruminant genomes including cattle, goat, and sheep (**Figure S2**). Subsequently, synteny analysis was applied to identify differences between the assembled genome and the domestic goat (*Capra hircus*) genome (**Figure S3**). 83.95% of two genome sequences could be 1:1 aligned, the average nuclear distance (percentage of different base pairs in the syntenic regions) was 7.18% (**Figure S4**). In addition, the density of different types of break points (edges of structural variation) are about 69.88 per Mb (**Table S3**). These results suggest that the reindeer genome assembly is of good level of contiguity and correctness.

### Genome annotation

To annotate the reindeer genome we initially used LTR_FINDER [11] and RepeatModeller (version 1.0.4; http://www.repeatmasker.org/RepeatModeler.html) to find repeats. Next, RepeatMasker (version 4.0.5) [12] was used (with -nolow -no_is -norna -parallel 1 parameters) to search for known and novel transposable elements (TE) by mapping sequences against the *de novo* repeat library and Repbase TE library (version 16.02) [13]. Subsequently, tandem repeats were annotated using Tandem Repeat Finder (version 4.07b; with 2 7 7 80 10 50 2000 -d -h parameters) [14]. In addition, we used RepeatProteinMask software [12] with -no LowSimple -*p* value 0.0001 parameters to identify TE-relevant proteins. The combined results indicate that repeat sequences cover about 1.07 Gb, accounting for 40.4% of the reindeer genome assembly (**Table S4**).

The rest of the reindeer genome assembly was annotated using both *de novo* and homology-based gene prediction approaches. For *de novo* gene prediction, we utilized SNAP (version 2006-07-28), GenScan [15], glimmerHMM and Augustus (version 2.5.5) [16] to analyze the repeat-masked genome. For homology-based predictions, sequences encoding homologous proteins of *Bos taurus* (Ensemble 87 release), *Ovis aries* (Ensemble 87 release) and *Homo sapiens* (Ensemble 87 release), were aligned to the reindeer genome using TblastN (version 2.2.26) with an (E)-value cutoff of 1 e-5. Genwise (version wise2.2.0) [17] was then used to annotate structures of the genes. The *de novo* and homology gene sets were merged to form a comprehensive, non-redundant gene set using EVidenceModeler software (EVM, version 1.1.1), which resulted in 21,555 protein-coding genes (**Table S5**). We then compared the reindeer genome with species which used in homology prediction, and there is no significant difference among the four species in gene length and exon length distribution (**Figure S5**).

Next, we searched the KEGG, TrEMBL and SwissProt databases for best matches to the protein sequences yielded by EVM software, using BLASTP (version 2.2.26) with an (E)-value cutoff of 1 e-5, and searched Pfam, PRINTS, ProDom and SMART databases for known motifs and domains in our sequences using InterProScan software (version 5.18-57.0). At least one function was assigned to 19,004 (88.17%) of the detected reindeer genes through these procedures (**Table S6**). The reads from short-insert length libraries then were mapped to the reindeer genome with BWA (version 0.7.12-r1039) [18], then called single nucleotide variant (SNV) by SAMtools (version 1.3.1) [19]. Finally, we performed SnpEff (version 4.30) [20] to identify the distribution of SNV in the reindeer genome (**Table S7**).

In addition, we predicted rRNA-coding sequences based on homology with human rRNAs using BLASTN with default parameters. To annotate miRNA and snRNA genes we searched the Rfam database (release 9.1) with Infernal (version 0.81), and annotated tRNAs using tRNAscan-SE (version 1.3.1) software with default parameters. The final results identified 159 rRNAs, 547 miRNAs, 1,339 snRNAs and 863 tRNAs (**Table S8**).

### Species-specific genes and phylogenetic relationship

We clustered the detected reindeer genes in families by using OrthoMCL [21] with an (E)-value cutoff of 1 e-5, and a Markov Chain Clustering with default inflation parameter in an all-to-all BLASTP analysis of entries for five species (*Homo sapiens, Equus caballus, Capra hircus, Bos taurus*, and *Rangifer tarandus*). The result showed that 335 gene families were specific to the reindeer (**Figure S6**). Moreover, we identified 7,505 single-copy gene families from these species and aligned coding sequences in the families using PRANK (version 3.8.31) [22]. Subsequently, 4D-sites (four-fold degenerated sites) were extracted to construct a phylogenetic tree by RAxML (version 7.2.8) [23] with GTR+G+I model. Finally, phylogenetic analysis using PAML MCMCtree (version 4.5) [24], calibrated with published timings of the divergence of the reference species (http://www.timetree.org/), indicated that *Rangifer tarandus*, *Bos taurus* and *Capra hircus* diverged from a common ancestor approximately 29.6 (25.4-31.7) Mya (**Figure S7**).

## Conclusion

In summary, we report the first sequencing, assembly and annotation of the reindeer genome, which will be useful in analysis of the genetic basis of the unique characteristics of reindeer, and broader studies on ruminants.

## Availability of supporting data

The raw data have been deposited in Genome Sequence Archive (GSA), under BIG Data Center, Beijing Institute Genomics (BIG), Chinese Academy of Science, with the project accession PRJCA000451.

## Abbreviations

Gb: giga base; bp: base pair; kb: kilo base; Mb: mega base; TE: transposable element; EVM: EVidenceModeler; BUSCO: benchmarking universal single-copy orthologs; FRC: feature-response curves; SNV: single nucleotide variant; Mya: million years ago

## Acknowledgements

This work was supported by the he Natural Science Foundation of China (No. 31501984) and Central Public-interest Scientific Institution Basal Research Fund (No. 1610342016026) to ZPL, and Talents Team Construction Fund of Northwestern Polytechnical University (NWPU) to QQ and WW. Special thanks to Nowbio Biotech Inc., Kunming, China for its remarkable work on DNA libraries constructions and sequencing.

## Competing interests

The authors declare that they have no competing interests.

## Authors’ contributions

ZPL collected the samples; ZSL, CL ZPL, YZ, KW and HB analyzed the data; ZSL, QQ and ZPL wrote the manuscript; GL, ZL, QQ and WW conceived the study.

## Figure legends

**Figure 1.**
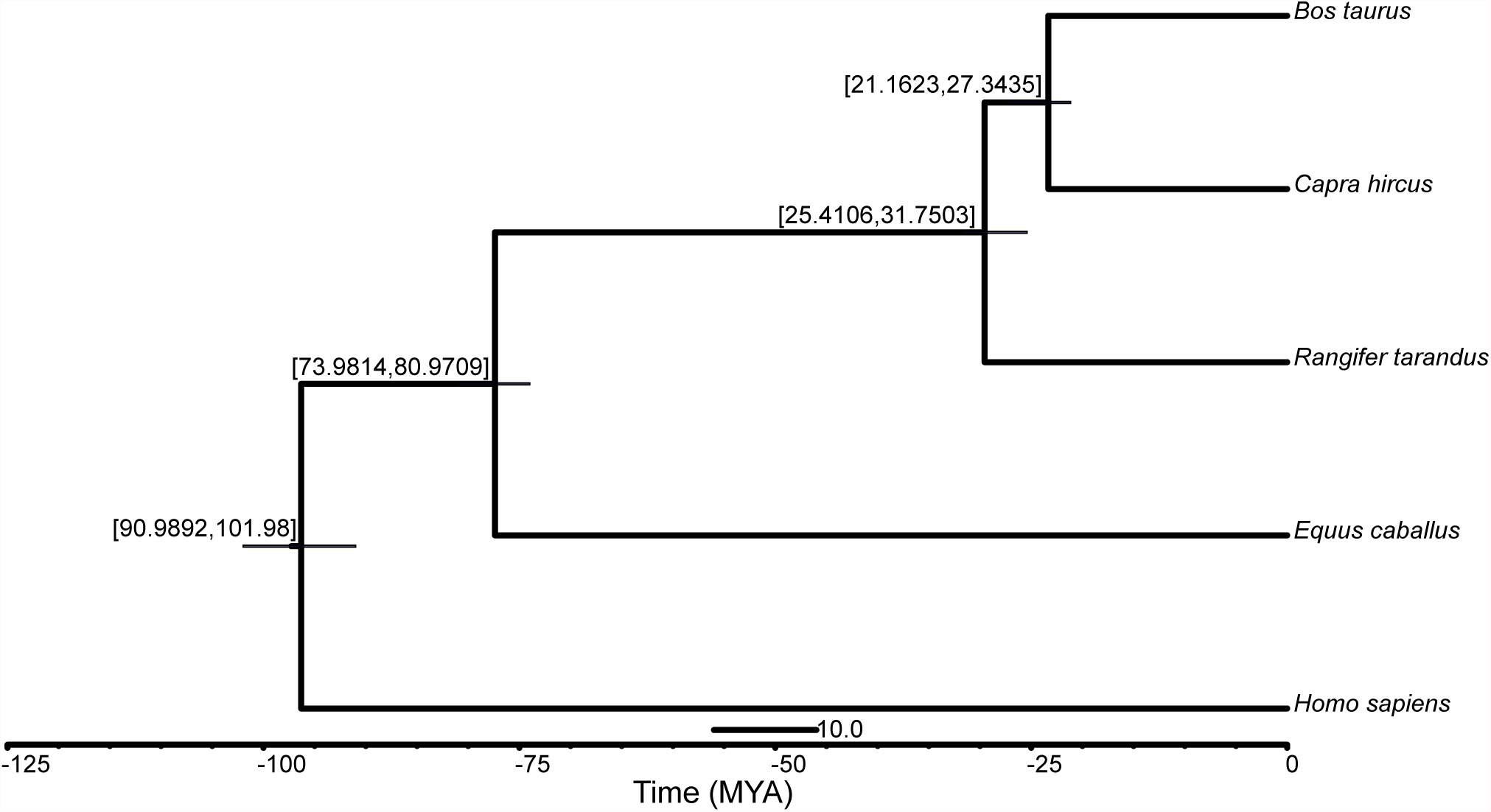
Phylogenetic relationships of *Rangier tarandus* and four species based on four-fold degenerated sites. Estimated divergence times are shown above the nodes. MYA, million years ago.

## References

1. Fernández MH and Vrba ES. A complete estimate of the phylogenetic relationships in ruminantia: a dated species-level supertree of the extant ruminants. Biological Reviews. 2005;80 2:269-302.

2. Hassanin A, Delsuc F, Ropiquet A, Hammer C, Jansen van Vuuren B, Matthee C, et al. Pattern and timing of diversification of Cetartiodactyla (*Mammalia*, *Laurasiatheria*), as revealed by a comprehensive analysis of mitochondrial genomes. C R Biol. 2012;335 1:32-50.

3. Young W. Park GFWH. Handbook of milk of non-bovine mammals. Wiley-Blackwell; 2006.

4. Luo R, Liu B, Xie Y, Li Z, Huang W, Yuan J, et al. SOAPdenovo2: an empirically improved memory-efficient short-read de novo assembler. GigaScience. 2012;1 1:1-6.

5. Li R, Fan W, Tian G, Zhu H, He L, Cai J, et al. The sequence and de novo assembly of the giant panda genome. Nature. 2010;463 7279:311-7.

6. Li R, Zhu H, Ruan J, Qian W, Fang X, Shi Z, et al. De novo assembly of human genomes with massively parallel short read sequencing. Genome Res. 2010;20 2:265-72.

7. Bickhart DM, Rosen BD, Koren S, Sayre BL, Hastie AR, Chan S, et al. Single-molecule sequencing and chromatin conformation capture enable de novo reference assembly of the domestic goat genome. Nat Genet. 2017;49 4:643-50.

8. Jiang Y, Xie M, Chen W, Talbot R, Maddox JF, Faraut T, et al. The sheep genome illuminates biology of the rumen and lipid metabolism. Science. 2014;344 6188:1168-73.

9. Simão FA, Waterhouse RM, Ioannidis P, Kriventseva EV and Zdobnov EM. BUSCO: assessing genome assembly and annotation completeness with single-copy orthologs. Bioinformatics. 2015;3119:3210-2.

10. Vezzi F, Narzisi G and Mishra B. Reevaluating assembly evaluations with feature response curves: GAGE and assemblathons. PLoS ONE. 2012;7 12:e52210.

11. Xu Z and Wang H. LTR_FINDER: an efficient tool for the prediction of full-length LTR retrotransposons. Nucleic Acids Res. 2007;35 suppl_2:W265-W8.

12. Tarailo-Graovac M and Chen N. Using RepeatMasker to identify repetitive elements in genomic sequences. Current Protocols in Bioinformatics. John Wiley & Sons, Inc.; 2009.

13. Jurka J, Kapitonov VV, Pavlicek A, Klonowski P, Kohany O and Walichiewicz J. Repbase update, a database of eukaryotic repetitive elements. Cytogenet Genome Res. 2005;110 1-4:462-7.

14. Benson G. Tandem repeats finder: a program to analyze DNA sequences. Nucleic Acids Res. 1999;27 2:573-80.

15. Burge C and Karlin S. Prediction of complete gene structures in human genomic DNA1. J Mol Biol. 1997;268 1:78-94.

16. Stanke M, Keller O, Gunduz I, Hayes A, Waack S and Morgenstern B. AUGUSTUS: ab initio prediction of alternative transcripts. Nucleic Acids Res. 2006;34 suppl_2:W435-W9.

17. Birney E, Clamp M and Durbin R. GeneWise and Genomewise. Genome Res. 2004;14 5:988-95.

18. Heng L. Aligning sequence reads, clone sequences and assembly contigs with BWA-MEM. arXiv. 2013;1303.3997

19. Li H, Handsaker B, Wysoker A, Fennell T, Ruan J, Homer N, et al. The sequence alignment/map format and SAMtools. Bioinformatics. 2009;25 16:2078-9.

20. Cingolani P, Platts A, Wang LL, Coon M, Nguyen T, Wang L, et al. A program for annotating and predicting the effects of single nucleotide polymorphisms, SnpEff: SNPs in the genome of Drosophila melanogaster strain w(1118); iso-2; iso-3. Fly (Austin). 2012;6 2:80-92.

21. Li L, Stoeckert CJ and Roos DS. OrthoMCL: Identification of Ortholog Groups for Eukaryotic Genomes. Genome Res. 2003;13 9:2178-89.

22. Löytynoja A and Goldman N. An algorithm for progressive multiple alignment of sequences with insertions. Proc Natl Acad Sci U S A. 2005;102 30:10557-62.

23. Stamatakis A. RAxML version 8: a tool for phylogenetic analysis and post-analysis of large phylogenies. Bioinformatics. 2014;30 9:1312-3.

24. Yang Z. PAML 4: Phylogenetic Analysis by Maximum Likelihood. Mol Biol Evol. 2007;24 8:1586-91.

